# Experimental parasite community perturbation reveals associations between Sin Nombre virus and gastrointestinal nematodes in a rodent reservoir host

**DOI:** 10.1101/2020.08.16.252288

**Authors:** Amy R. Sweeny, Courtney A. Thomason, Edwin A. Carbajal, Christina B. Hansen, Andrea L. Graham, Amy B. Pedersen

## Abstract

Individuals are often co-infected with several parasite species, yet measuring within-host interactions remains difficult in the wild. Consequently, such interactions’ impacts on host fitness and epidemiology are often unknown. We used anthelmintic drugs to experimentally reduce nematode infection and measured the effects on both nematodes and the important zoonosis Sin Nombre virus (SNV) in its primary wild reservoir (*Peromyscus spp*.). Treatment significantly reduced nematode infection, but increased SNV seroprevalence. Furthermore, mice that were co-infected with both nematodes and SNV were in better condition and survived as much as four times longer than mice infected with either parasite alone. These results highlight the importance of investigating multiple parasites for understanding interindividual variation and epidemiological dynamics in reservoir populations with zoonotic transmission potential.

## (1) Introduction

Co-infection with both micr0parasites and macroparasites is common in the wild [1,2]. Interactions among parasites co-habiting a host can occur through multiple mechanisms including bottom-up (e.g. resource competition) or top-down (e.g. immune-mediated) processes [3-5]. These interactions can alter both host and parasite fitness [1,2,6,7], e.g. increasing parasite burdens for a co-infecting species [8,9], worsening disease pathology [6], altering transmission rates [10], and ultimately influencing efficacy of disease control strategies [11].

Disease ecologists commonly assess the consequences of infection in a wild host by removing a target parasite group using antiparasite drug treatments [12], but monitoring the non-target parasite community response is rarer. Some studies have used perturbation experiments to determine the strength and direction of within-host parasite interactions by measuring the response of non-targeted parasite species after treatment [8,9,13]. For example, in African buffalo (*Syncerus caffer*), animals treated to remove nematodes were nine times more likely to survive co-infection with the bacterium *Mycobacterium tuberculosis* [13]. In contrast, the removal of nematodes in wild rodents has been shown to increase coccidian microparasite infection, possibly through a competitive release response [8,9]. These studies show that ignoring the broader parasite community may crucially underestimate the occurrence and importance of within-host interactions and suggest that experimental approaches can provide mechanistic insight into these relationships [14,15]. Co-infection poses substantial public health risks, especially for parasites with zoonotic potential or that severely impair the immune system – as with HIV and the re-emergence of drug-resistant tuberculosis [16].

Small mammals disproportionately serve as reservoir species for zoonotic diseases [17-20] and are ideal, tractable systems for experimental studies. Hantavirus Pulmonary Syndrome (HPS) represents an important zoonotic disease caused by Sin Nombre virus (SNV), endemic in deer mice (*Peromyscus maniculatus*) and white-footed mice (*P*. *leucopus*) [21,22]. Hantavirus infection can reduce wild rodent fitness [22,23] and often co-occurs with other endemic parasites [24,25]. Here we experimentally perturbed the taxonomically diverse parasite communities of deer and white-footed mice, the primary wild reservoirs of SNV [21]. Nematodes represent a keystone parasite in within-host communities because they can interact with other parasites through the host immune system or through direct competition for resources in the gastrointestinal (GI) tract [4,26]. Previous work in this system found that GI nematode infections were common and interacted with other co-infecting GI and ectoparasites [27,28]. We used anthelmintic treatment to remove nematodes and monitored downstream effects on SNV infection and host fitness. We show that removal of nematode infections increases subsequent probability of SNV seroconversion and that co-infection with nematodes and SNV conveys condition and survival benefits within this population.

## (2) Methods

Field experiments were conducted at the Mountain Lake Biological Station (MLBS) in southwest Virginia, where populations of deer and white-footed mice have been monitored for decades [29], and the parasite community is well-characterised [9,27]. Live-trapping took place from May/June to August in two temporal replicates (2010-2011) on three spatially-matched 0.5 ha grid replicates (8 x 8 trap arrays; 10m spacing). Each spatially-matched grid set was trapped for three consecutive nights every two weeks. In each temporal replicate, experimental design included randomized anthelmintic treatment (administered at first capture and repeated fortnightly) with a weight-adjusted oral dose of Ivermectin (5 mg/kg; Eqvalan, Merial, USA) or control (5% sucrose solution). See Electronic Supplementary Material (ESM) for additional details.

At first capture, individuals were ear-tagged and their species identified using morphological characteristics [30]. At each capture, morphometric data including age, sex, weight, length, and reproductive condition were recorded. Fecal and blood samples were also collected at each fortnightly capture. The presence/absence and number of eggs/gram faeces (a common proxy of infection intensity; EPG) for nematodes species were quantified using a salt flotation technique [31]. Nematode species were aggregated for analysis because drug treatment is at the group (nematode) level. Blood samples were screened for SNV antibodies using standard enzyme-linked immunosorbent assay (ELISA) protocols and reagents from the U.S. Centers for Disease Control and Prevention [32,33]. ELISA results were used to assign infection status based on seropositivity (presence/absence: threshold of 3SD > negative control) and for positive samples the adjusted optical density (OD) relative to a negative control (CDC # 703226) was used to estimate concentration of antibodies for statistical analyses. Additional details in ESM.

All statistical analyses were conducted in R v3.6.0 (R Core Team 2013). We first investigated factors driving natural SNV and nematode infections prior to experimental perturbations by fitting generalised linear models (GLMs) using the package ‘glmmTMB’ to SNV (both presence/absence and antibody concentration) or nematodes (both presence/absence and intensity (egg/gram feces)) for all first capture events. Models were fit with with binomial (logit link; SNV & nematode presence/absence), gaussian (SNV OD-positive only) or negative binomial (log link; nematode intensity, infected only) error distributions. We included the following fixed effects: year (Factor: 2011/2012), julian date of capture (continuous, scaled to mean = 0 / SD = 1), sex (Factor: male/female), age (Factor: sub-adult/adult), species (Factor: *P*. *leucopus*/*P*. *maniculatus*), and body weight (grams, continuous).

We then explicitly tested the relationship between SNV and GI nematodes by fitting generalized linear mixed-effects models (GLMM) to the same response variables detailed above, using data from all captures and including additional fixed effects of treatment (2-level factor: ivermectin treated/ control), nematode infection status (2-level factor: present/absent), and treatment*timepoint (2-level factor: pre-/post-treatment). Addition model details in ESM.

Finally, we investigated the effects of drug treatment and SNV-nematode co-infection on host body weight as a proxy of condition and recapture duration (number of days known alive) as a proxy for survival using a GLM with gaussian and negative binomial error distributions respectively. The following fixed effects were included for both models: year, sex, age, species, reproductive status, treatment (all as described above), and infection status (Factor, 4-level: None, SNV only, Nematode only, Co-infected). Condition models included an additional random effect of individual ID, while survival models included additional fixed effects of weight and trap session across both years (continuous, 1-11) to account for skewed observation times. Grid (6-level factor) was included as a random effect in all models to account for variation across spatial replicates.

## (3) Results

### (a) Parasite dynamics

539 individuals were captured in the field experiments (2010: n=239; 2011: n=300). Prior to anthelmintic treatment, SNV prevalence was 8.6% and GI nematode prevalence was 26.8%. Additionally, prior to treatment, mouse sex, body weight, and time of season were the primary determinants of both SNV infection probability and antibody concentration (Table 1). Males (Sex, Male: Infection probability - β = 0.89, SE = 0.39, p = 0.022; Titer - β = 0.11, SE = 0.06, p = 0.09) and larger mice (Weight (g): Infection probability - β = 0.11, SE = 0.05, p = 0.015; Titer - β = 0.02, SE = 0.01, p = 0.048) were more likely to be infected with SNV, while infection probability declined later in the summer (Julian date (scaled): Infection probability - β = −0.51, SE = 0.21, p = 0.013; Titer - β = −0.12, SE = 0.03, p < 0.001). There were no significant predictors of nematode infection probability at first capture (Table 1), and only time of season was a significant predictor of nematode infection intensity (Julian date (scaled): β = 0.90, SE = 0.26, p < 0.001), where EPG increased over the course of the summer.

**Table 1.**
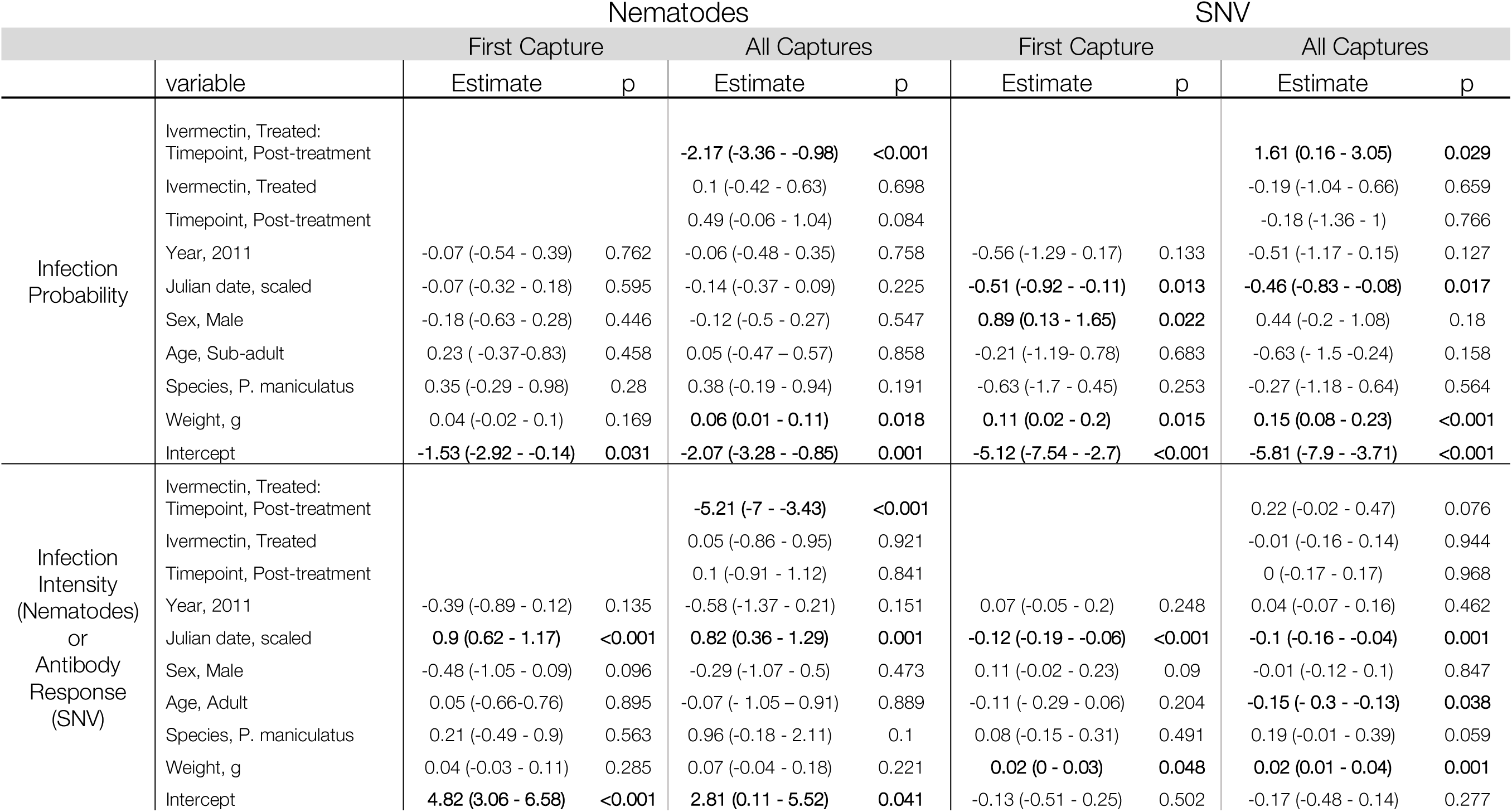
Model output for Nematode and SNV dynamics at first capture and all captures of the experiment.

### (b) Drug treatment effects on parasite infection & host fitness

Anthelmintic treatment significantly reduced nematode infection probability (77.0% reduction) and intensity (89.8% reduction) (Ivermectin, Treated:Timepoint, Post-Treatment: Probability - β = −2.17, SE = 0.61, p < 0.001; Intensity - β = −5.21, SE = 0.91, p < 0.001) (Figure 1A-B). In contrast, SNV infection probability increased following anthelmintic treatment (54.5% increase, Probability - β = 1.61, SE = 0.74, p = 0.029). SNV titer was also greater following anthelmintic treatment, however this increase was not significant (Titer - β = 0.22, SE = 0.13, p = 0.076). Body weight was an additional predictors of nematode infection probability, where larger mice were more likely to be infected (Table 1). As with the parasite dynamics models, capture date was a predictor of nematode infection intensity, and body weight and capture date were significant predictors of SNV infection probability and antibody concentration (Table 1). Finally, sub-adult mice had higher antibody response to SNV compared to adult mice (Table 1).

**Figure 1.**
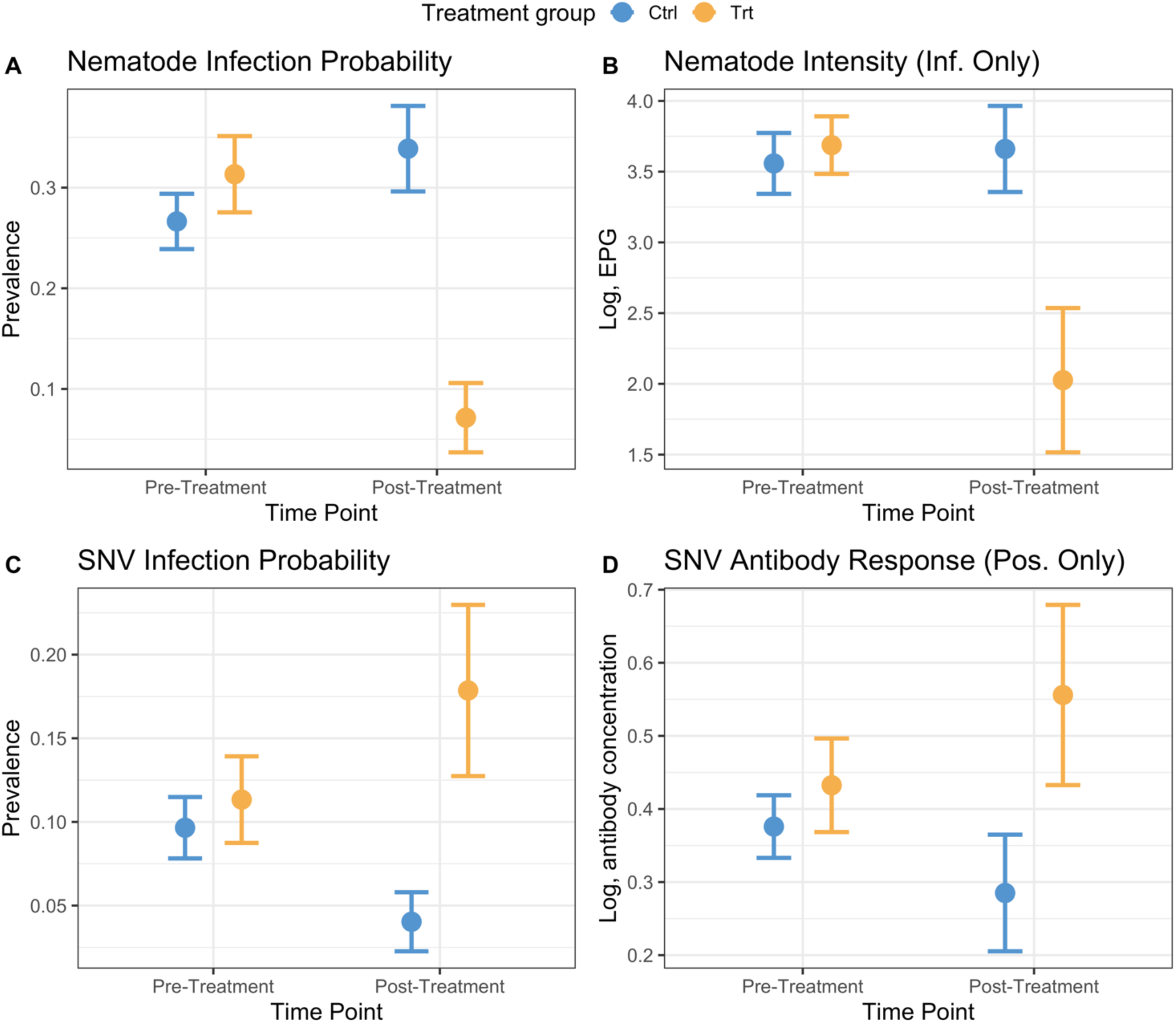
Anthelmintic treatment impacts nematode (A-B) and SNV (C-D) infection dynamics. Plots represent raw data for pre- or post-treatment groups. Treated mice (orange) had lower nematode infection probability (A) and intensity (B), but higher SNV infection probability (C) and antibody response (D). Points represent mean ± SE.

We found positive effects of co-infection with SNV and nematodes on both host body weight as a proxy of host condition (Weight, g: β = 3.20, SE = 0.83, p < 0.001) and recapture duration as a proxy for survival (Observation length, days: β = 1.52, SE = 0.26, p < 0.001), where co-infected individuals on average were 3 g (20%) heavier and observed for 4x longer than singly-infected individuals (Figure 2). In weight models, variation of weight with age was accounted for by including host age as a fixed effect (Age Class, Adult: β = 5.04, SE = 0.35, p < 0.001). We found additional effects of sex (Sex, Male: β = −1.10, SE = 0.34, p = 0.001) and reproductive status (Repro status, active: β = 1.61, SE = 0.31, p < 0.001) on body weight. In survival models, time into the experiment to first capture was accounted for by including trap session as a fixed effect (β = −0.14, SE = 0.058, p = 0.019), where the later the date of first capture the shorter the observation length. Adults had slightly higher observation lengths compared to sub-adults, but this effect was marginal (β = 0.35, SE = 0.19, p = 0.059).

**Figure 2.**
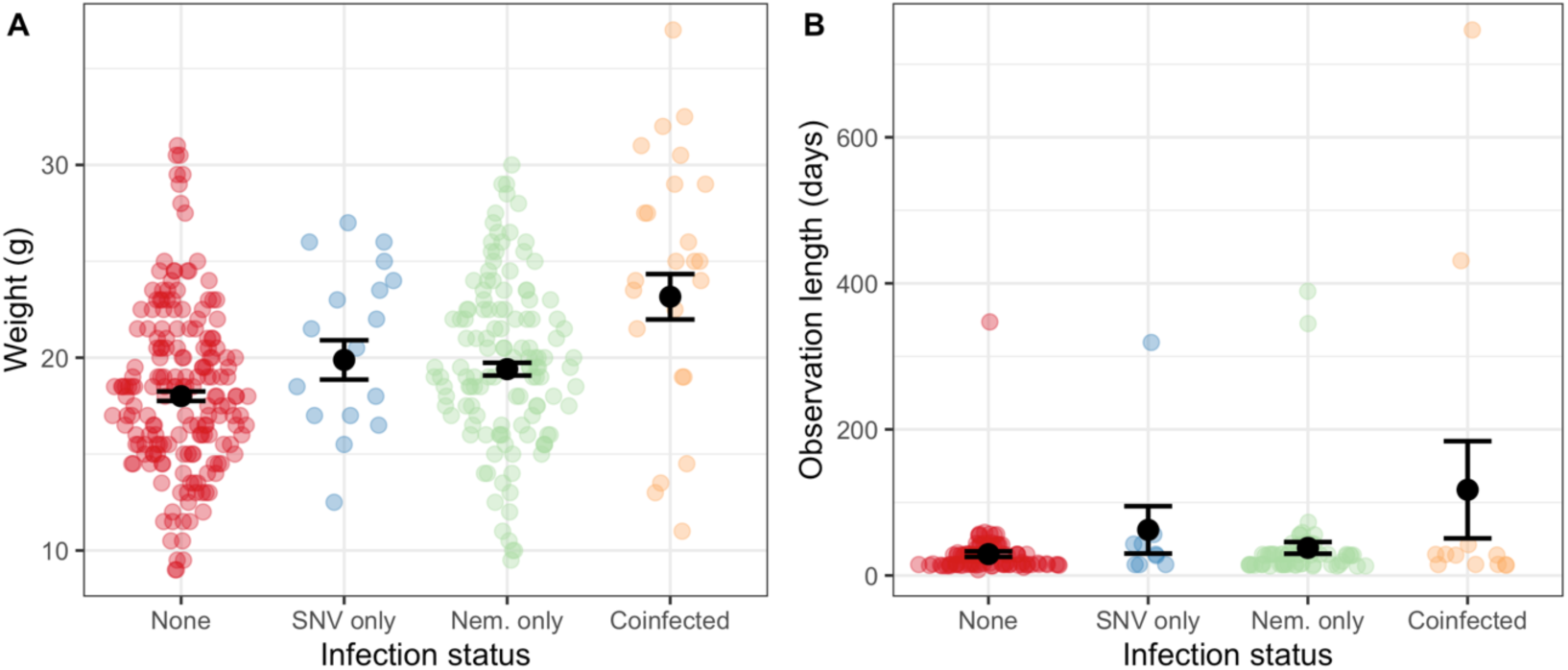
Co-infection effects on (A) host condition (weight, g) and (B) host survival (days observed). Sina plots represent raw data distributions for infection groups: never infected, infected with either SNV or nematodes, or co-infected. Points represent mean ± SE.

## (4) Discussion

Efforts to understand the risk of emerging infectious disease from wildlife reservoirs commonly focus on anthropogenic or environmental factors that influence contact at the human-wildlife interface [34-36], while interindividual variation in susceptibility and transmission potential within reservoir hosts remains under-studied [37,38]. Within-host interactions among parasites can shape infection risk and fundamentally change pathogen virulence and transmission potential [14], but the influence of co-infection on zoonotic potential is still poorly understood. Here, we show that the loss of important nematode parasites drove increased prevalence of a zoonotic virus, demonstrating that coinfecting parasites could be an important mediating factor in determining wild animal populations’ zoonotic potential. This finding supports the idea that parasite diversity loss could result in increased zoonotic outbreaks [39] and that parasite conservation effects may be a valuable strategy in zoonotic disease control [40].

Mice that were co-infected with SNV and GI nematodes had higher body weight and were observed for longer than uninfected or singly-infected individuals, which could alter disease dynamics by modifying infected individuals’ transmission potential. For example, anthelminthic treatment in African buffalo (*Syncerus caffer*) decreased mortality from bovine tuberculosis (BTB), resulting in an eightfold increase in BTB the reproductive number within the population [13]. Although hantaviruses are not considered to cause much pathology in rodents, evidence from two field studies of *Peromyscus spp*. suggests that they can result in some associated mortality [22,23]. The enhanced condition and lifespan observed here may represent an unexpected benefit of nematode co-infection for wild rodents infected with SNV. Our analysis uses observed time alive from the trapping data as a proxy for survival, so we cannot explicitly rule out dispersal as a confounding event; however, previous work suggests that SNV does not influence dispersal in deer mice [41]. Furthermore, it should be noted that a small number of mice (N=10) survived for multiple years in this study and contribute disproportionately to observation length within the co-infected group, and direction of causality cannot be assumed with this sample size. Further work is therefore needed to explicitly test these effects.

Our observations imply that reducing nematode load creates beneficial conditions for SNV infection, potentially by altering the within-host immune environment. The immune response to nematodes is typically dominated by a combination of T-helper cell 2 (Th2) and T-helper cell regulatory (Treg) immune responses [42]. These responses include a suite of Th2-related cytokines, which are important mediators of inflammatory responses of the T helper cell 1 (Th1) arm of the immune system [43], and are better suited to minimize damage to the host rather than directly clear parasites, resulting in chronic infections [44]. Hantaviruses are likewise chronic in rodents and use a distinct mechanism to achieve immune evasion to persist and replicate in the absence of overt disease, whereby the virus may directly mediate suppression of Th1 responses via structural and non-structural proteins [45]. Antibody responses develop two weeks post hantavirus infection and can remain detectable throughout a rodent’s life, providing protection, and in some cases elimination. It is therefore possible that our results represent a reversal of nematode-induced immunosuppression following anthelmintic treatment. Given the infection status of SNV was determined by ELISA assays, it is also possible that these results reveal increased detection probability of SNV due to the higher magnitude of response in the absence of nematode infection. Alternatively, if chronic nematode infection imposes energetic costs, removing these parasites could result in greater host movement and sociality, driving greater SNV exposure [46,47]. Regardless of underlying mechanisms, these observed nematode-SNV interactions confirm that distantly-related parasites can be mechanistically linked, and studies that do not consider co-infection may be missing an important source of variation in disease ecology [3,14,48].

## Acknowledgments

We thank A. Greenlee, K. Grannum, D. Kone, O. Yin, C. Rueb, and T. Marion for their fieldwork assistance, Greg Albery for insightful comments, and Mountain Lake Biological Station of University of Virginia for their support.

## Author Contributions

This study was conceived of by CAT, ALG, and ABP. Fieldwork was carried out by CAT, ARS, and EAC. Lab assays were carried out by CAT, ARS, EAC, and CAH. ARS and CAT analysed the data and prepared first draft of the manuscript. All authors contributed to the writing of the final manuscript.

## Data Accessibility Statement

Data will be available on Dryad Digital Repository upon acceptance.

## Electronic Supplementary Material

### Field Experiment & Data Collection

Mountain Lake Biological Station is located in an oak-maple forest the Appalachian Mountains of southwestern Virginia (37°10’N, 80°20’W, 1200 m). Trapping nights consisted of baiting Sherman live traps (3” × 3.5” × 9”, H. B. Sherman Traps, Tallahassee, FL) with crimped oats and cotton bedding on cool (<55°F) nights and checking for captures early the next morning (Figure S1). Animal processing and sampling was carried out in accordance with the Texas Tech University IACUC guidelines under protocol numbers 09013-4 and the Virginia Department of Game and Inland Fisheries small mammal trapping permit # 041919.

**Figure S1.**
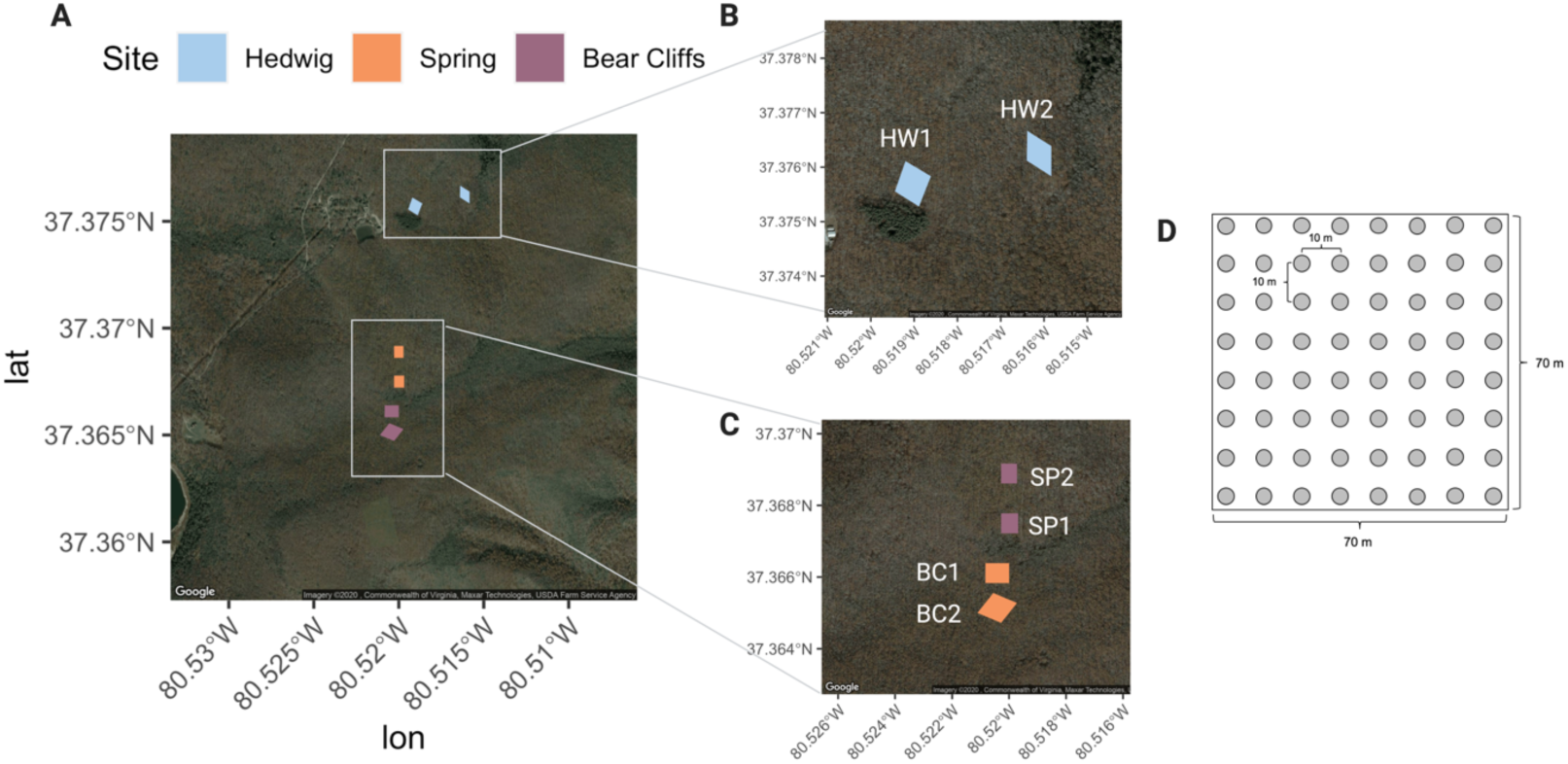
Grid layout for trapping sites included in field experiment analysis. A. Arrangement of all 3 trapping sites (6 grids) surround Mountain Lake Biological Station. B. Grid location of site Hedwig. C. Grid locations of sites Spring & Bear Cliffs. D. Schematic of trapping location layout within each grid.

In 2010, treatment was assigned at the population-level, where one of each pair of spatial replicate grids was assigned to anthelmintic treatment (all animals were treated) and the other assigned to control (5% sucrose solution). In 2011, treatment was assigned at the individual level for 4/6 grids and captured mice were assigned to one of four treatment groups: (i) ivermectin, (ii) fipronil, (iii) ivermectin + fipronil, or (iv) control. The remaining two grids were designated as control. Fipronil, a common insecticide that was administered topically, was used to reduce arthropod vectors (e.g. fleas) that can transmit additional microparasites as part of addition investigation into parasite community dynamics. However, we found no effect of fipronil treatment on either nematodes or SNV so these treatments were grouped as either received ivermectin (treated) or no treatment (control) for subsequent analyses.

Morphometric data collection was carried out as follows. Length (body, tail, hindfoot, ear) measurements were recorded, and *Peromyscus maniculatus* was distinguished from *P*. *leucopus* based on tail length exceeding body length, sharply bicoloured tail and a hair tuft at the end of the tail. Age (juvenile, sub-adult or adult) was assessed for both species by pelage color and molt patterns. Sex was determined by the urogenital distance. Mice were considered reproductively active if the females had a perforate vagina, were pregnant, or lactating; or testes were descended in males.

Retro-orbital bleeds were carried out under short-term anesthesia (Isoflurane, ∼150 µl). Faecal samples were collected from individual traps at the end of each day, and traps were subsequently washed with hospital-grade detergent before re-use.

### Laboratory assays

Fecal samples were weighed and stored in 10% formalin prior to parasite quantification. Blood samples were centrifuged to separate plasma and stored at −80°C until serology assays. SNV serology followed the standard CDC protocol, as follows. ELISA microtiter plates (Nunc Star Well Maxisorb, Thermo Scientific 441653) were coated overnight at 4°C with 100 μL per well of recombinant affinity purified SNV antigen (CDC Lot No. SPR569, 1:2000 dilution in phosphate buffered saline (PBS), 0.01 M, pH 7.4) on half of each plate and a control antigen (containing components of the cell lines used to produce the SNV antigen; CDC Lot No. SPR568, 1:2000 dilution in PBS) on the other half. Plates were then washed three times with PBS supplemented with 0.1% Tween-20 (PBST, pH 7.4). Serum samples were diluted 1:100 in ELISA diluent (PBST supplemented with 5% skim milk, pH 7.4) and added to wells. Positive control serum (anti-SNV NC antigen HMAF; CDC #703142) was used at an initial dilution of 1:1000, and negative control serum (CDC # 703226) at 1:100, both in ELISA diluent. Positive and negative controls were included on each plate for each antigen.

After incubation of samples and controls at 37°C for 60 minutes, wells were washed three times with PBST. Peroxidase-labeled anti-*Rattus norvegicus* conjugate (Kirkegaard and Perry Laboratories, Cat No. 14-16-06, diluted 1:2500 in ELISA diluent) and peroxidase-labeled anti-*Peromyscus leucopus* IgG secondary antibody (Kirkegaard and Perry Laboratories, Cat No. 14-33-06, diluted in 1:2500 in ELISA diluent) were then added to all wells and incubated for 60 minutes at 37°C. Plates were washed three times and then ABTS peroxidase substrate (Kirkegaard and Perry Laboratories, Cat. Nos. 506400 & 506500, combined 1:1) was added (100 μL per well). Plates were then incubated for 30 minutes at 37°C. Spectrophotometric data were collected at 405nm on a Multiskan GO.

Serological testing was done in two stages. First, all samples were screened for seropositivity in quadruplicate wells. For each serum sample, we subtracted the mean OD values of the wells with control antigen from the mean OD values of the wells with the SNV antigen, to give a “net” positive adjusted OD value. Results were considered positive if this value was greater than 3 standard deviations above the adjusted OD405 (cut-off value) for the negative control samples.

### Statistical analysis

We use a timepoint:treatment interaction here to test group differences. Individual ID was initially included in the treatment models as a random effect, but dropped given the very high percentage (>70%) of individuals which had only single captures, resulting in non-convergence of models estimating interindividual variation (Figure S2).

**Figure S2.**
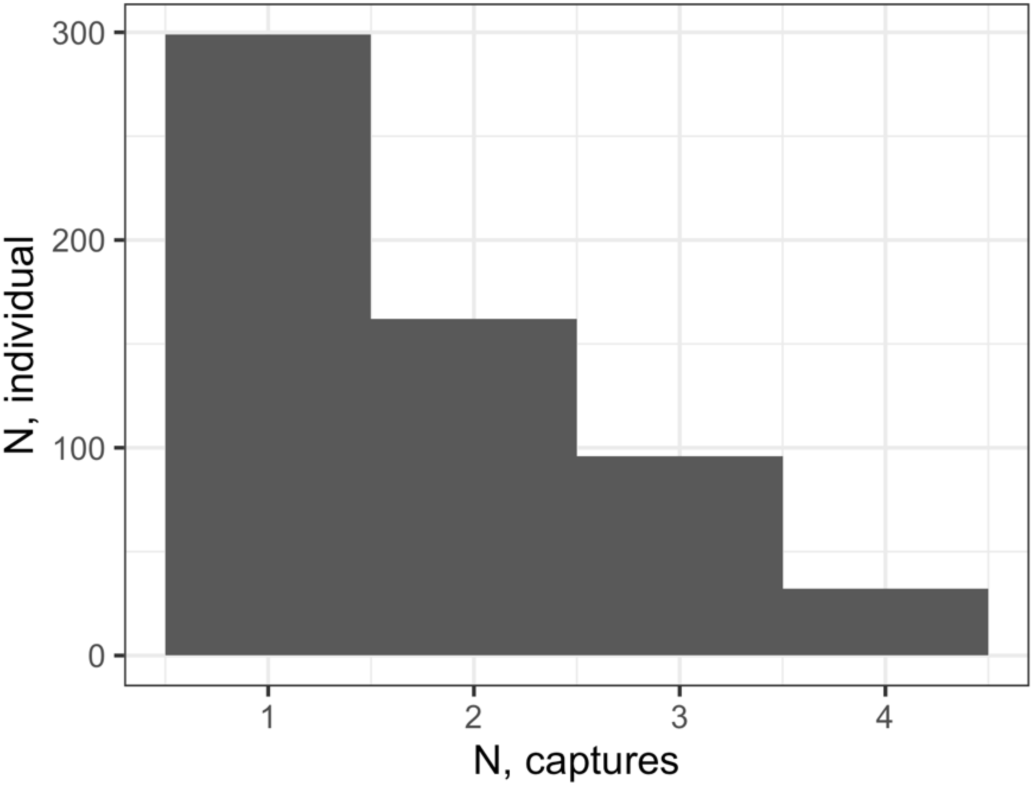
Histogram of capture frequency for mice included in field experiment. 299 (71.2%) mice were captured only once and 81 (19.3%) had two captures, with very few (32 and 8, respectively) having beyond 2 (3 or 4) captures.

